# RepeatModeler2: automated genomic discovery of transposable element families

**DOI:** 10.1101/856591

**Authors:** Jullien M. Flynn, Robert Hubley, Clément Goubert, Jeb Rosen, Andrew G. Clark, Cédric Feschotte, Arian F. Smit

**Affiliations:** Department of Molecular Biology and Genetics, Cornell University, Ithaca, NY 14853, USA; Institute for Systems Biology, Seattle, WA 98109, USA

## Abstract

The accelerating pace of genome sequencing throughout the tree of life is driving the need for improved unsupervised annotation of genome components such as transposable elements (TEs). Because the types and sequences of TEs are highly variable across species, automated TE discovery and annotation are challenging and time-consuming tasks. A critical first step is the de novo identification and accurate compilation of sequence models representing all the unique TE families dispersed in the genome. Here we introduce RepeatModeler2, a new pipeline that greatly facilitates this process. This new program brings substantial improvements over the original version of RepeatModeler, one of the most widely used tools for TE discovery. In particular, this version incorporates a module for structural discovery of complete LTR retroelements, which are widespread in eukaryotic genomes but recalcitrant to automated identification because of their size and sequence complexity. We benchmarked RepeatModeler2 on three model species with diverse TE landscapes and high-quality, manually curated TE libraries: *Drosophila melanogaster* (fruit fly), *Danio rerio* (zebrafish), and *Oryza sativa* (rice). In these three species, RepeatModeler2 identified approximately three times more consensus sequences matching with >95% sequence identity and sequence coverage to the manually curated sequences than the original RepeatModeler. As expected, the greatest improvement is for LTR retroelements. The program had an extremely low false positive rate when applied to simulated genomes devoid of TEs. Thus, RepeatModeler2 represents a valuable addition to the genome annotation toolkit that will enhance the identification and study of TEs in eukaryotic genome sequences. RepeatModeler2 is available as source code or a containerized package under an open license (https://github.com/Dfam-consortium/RepeatModeler, https://github.com/Dfam-consortium/TETools).

**Significance:** Genome sequences are being produced for more and more eukaryotic species. The bulk of these genomes is composed of parasitic, self-mobilizing transposable elements (TEs) that play important roles in organismal evolution. Thus there is a pressing need for developing software that can accurately identify the diverse set of TEs dispersed in genome sequences. Here we introduce RepeatModeler2, an easy-to-use package for the curation of reference TE libraries which can be applied to any eukaryotic species. Through several major improvements over the previous version, RepeatModeler2 is able to produce libraries that recapitulate the known composition of three model species with some of the most complex TE landscapes. Thus RepeatModeler2 will greatly enhance the discovery and annotation of TEs in genome sequences.

## Introduction

Most eukaryotic genomes contain a large number of interspersed repeats that by and large represent copies of transposable elements (TEs) at varying stages of evolutionary decay (Smit 1999; Consortium and International Human Genome Sequencing Consortium 2001; Jurka *et al*. 2007; Huang *et al*. 2012; Bourque *et al*. 2018). TEs are genomic sequences capable of mobilization and replication, generating complex patterns of repeats that account for up to 85% of eukaryotic genome content (International Wheat Genome Sequencing Consortium (IWGSC) *et al*. 2018). Different organisms have diverse TE landscapes, including a wide range of abundances, activity levels, and sequence degradation levels (Hua-Van *et al*. 2005; Smit 2012). As mutagens and major contributors to the organization, rearrangement, and regulation of the genome, TEs have had a profound impact on organismal evolution (reviewed in (Bourque *et al*. 2018)). Our understanding of the biological impact of TEs has grown steadily through the study of both model and non-model organisms from which whole genome sequences can now be routinely assembled. With each new species sequenced comes the challenge of identifying its unique set of TE families, which remains a tedious and largely manual endeavor. Yet, the accurate identification of TEs and other repeats is prerequisite to nearly all other genomic analysis, including the annotation of genes (Yandell and Ence 2012).

What makes TEs so potent in remodeling the genome but also challenging to annotate is their diversity in structures and sequences, which greatly vary across species (Huang *et al*. 2012; Bourque *et al*. 2018). There are two major classes of TEs (reviewed in (Finnegan 1989; Wicker *et al*. 2007; Piégu *et al*. 2015); https://www.dfam.org/classification): class I retroelements replicate and transpose via an RNA intermediate; while class II elements (or DNA transposons) are mobilized via a DNA intermediate. Class I elements include long and short interspersed elements (LINEs, SINEs) and long terminal repeat (LTR) retrotransposons. The most common class II transposons are TIR (terminal inverted repeats) elements, which transpose via a “cut-and-paste” mechanism (Feschotte and Pritham 2007). But other class II elements, such as *Helitrons*, also use replicative mechanisms (Thomas and Pritham 2015; Grabundzija *et al*. 2018). Within each class, TE sequences are extremely diverse and evolve rapidly (Wicker *et al*. 2007; Arkhipova 2017; Bourque *et al*. 2018). Additionally, once integrated in the host genome each element is subject to mutations, such as point mutations, and a vast array of rearrangements, including internal deletions, truncations, and nested insertions. The vast sequence diversity of TEs combined with the complexity of mutations that occur after insertion makes automated TE identification and classification a daunting task.

The most elementary level of classification of TEs is the family, which designates interspersed genomic copies derived from the amplification of an ancestral progenitor sequence (Wicker et al. 2007). Each TE family can be represented by a consensus sequence approximating that of the ancestral progenitor. Such consensus sequence can be recreated from a multiple alignment of individual genomic copies (or “seeds”) from which each ancestral nucleotide can be inferred based on a majority-rule along the alignment. Similarly, the seed alignment may be used to generate a profile Hidden Markov Model for each family. Consensus TE sequences and HMMs are used for many downstream applications in the study and annotation of genomes. Notably they are used to annotate or “mask” the genome using RepeatMasker, which is a prerequisite for gene annotation (Yandell and Ence 2012). Consensus sequences are generally stored in widely used databases such as Repbase (Bao *et al*. 2015) or along with seed alignments and HMMs in Dfam (Hubley et al. 2016). Seed alignments and accurate sequence models are critical for reconstructing the evolutionary history of TEs and are used for a variety of biological studies including the study of TE invasions and regulation (e.g. (Kofler et al. 2015)). Years of manual curation have resulted in high quality consensus libraries for a limited set of species, mostly model organisms (Lerat *et al*. 2003; Hubley *et al*. 2016; Stitzer *et al*. 2019).

The number of whole-genome assemblies for species throughout the tree of life continues to grow at a very fast rate, and efforts are underway to produce thousands more (Koepfli *et al*. 2015; Lewin *et al*. 2018). Long-read sequencing technologies are improving the quality of genome assemblies, especially in highly repetitive regions (e.g. (Chang and Larracuente 2019)). These developments bring a pressing need to improve tools that automate the discovery and annotation of TEs. Although there are dozens of tools that tackle one aspect of *de novo* identification or one class of TE (Saha *et al*. 2008a; b), there are very few easy-to-use programs that can produce a comprehensive library of TE family seed alignments and consensus sequences from a genome assembly.

RepeatModeler was released in 2008 by Hubley and Smit and is one of the most widely used TE discovery tools (cited 1462 times in publications as of 11/21/2019). RepeatModeler constructs seed alignments and consensus sequences for genome-wide repeat families *de novo*. However, the original version of RepeatModeler, like other existing TE-discovery software, falls short of producing a complete, non-redundant library of full-length consensus sequences. The most problematic issue is the representation of what should be a unique contiguous consensus sequence for a given TE family into many fragmented and partially redundant sequences in the output library. This issue, in turn, can hamper the classification of the TE families, inflates the number of actual TE families in the genome, and confounds genome annotation and downstream analyses. LTR retroelements are particularly recalcitrant to automated TE identification because of their size (up to 20 Kbp) and complexity in sequence and organization, which is driven in part by their ability to recombine within and between families (Vargiu *et al*. 2016). Yet these elements are widespread and often extremely abundant and diverse in eukaryotic genomes. For instance, the maize reference genome harbors >100,000 LTR elements falling into ~20,000 distinct families accounting for about half of the genomic DNA (Jiao *et al*. 2017).

To address these issues we developed a new version of RepeatModeler. Notably, we integrated an optional module dedicated to the identification of LTR elements in the genome through their structural characteristics (Ou and Jiang 2018; Ellinghaus *et al*. 2008). By benchmarking on three diverse model species, we demonstrate that RepeatModeler2 is a substantial improvement over the previous version both in terms of detection sensitivity and consensus sequence quality. The open-source package is designed to run on a single, multi-processor computer and is available as a source distribution or Docker/Singularity container for easier installation. (https://github.com/Dfam-consortium/RepeatModeler).

## Methods

### RepeatModeler2 overview

RepeatModeler is a pipeline for automated *de novo* identification of TEs that employs two distinct discovery algorithms, RepeatScout (Price *et al*. 2005) and RECON (Bao 2002), followed by consensus building and classification steps. In addition, RepeatModeler2 now includes the LTRharvest (Ellinghaus *et al*. 2008) and LTR_retriever (Ou and Jiang 2018) tools. Our tool takes advantage of the unique strengths of each approach as well as providing a tractable solution to analyzing large datasets such as whole genome assemblies. For instance, RepeatScout uses high frequency sequence word counts to identify interspersed repeat seeds (short regions of putative homology) and then performs an iterative extension of a multiple alignment around the aligned seeds, similar to the seed and extend phase of the BLAST pairwise alignment algorithm. While RepeatScout’s implementation of this algorithm is fast, the program input is currently limited to ~1 Gbp and the alignment scoring system (+1/-1, and non-affine gap penalty) limits the divergence of discovered families. Despite these limitations, RepeatScout serves well as a fast method to discover the youngest and most abundant families given a small sequence sample from a genome. RECON, on the other hand, provides sophisticated and TE biology-aware clustering and relationship determination approaches to generate TE families from exhaustive inter-genome alignments. RECON’s approach requires a computationally intensive but sensitive alignment (sophisticated scoring matrices, and affine gap penalties) and detects older TE families quite well.

In order to comprehensively identify TE families in a genome we chose to employ a sampling and iterative (sample, mask, identify) search strategy (Figure 1). We begin by supplying RepeatScout with a random 40 Mbp sample of the genome to quickly identify young and abundant families. In each successive round we mask a new genomic sample using all previously discovered TE families to avoid re-discovery and allowing for larger successive sample sizes as the computational burden of self-comparison is reduced on pre-masked sequence. The second and subsequent rounds all employ the RECON approach starting with a 3 Mbp sample (without replacement), tripling the sample size between rounds and continuing until a sample size maximum or round limit is reached (default: 243 Mbp, or 5 rounds). For an average mammalian genome of 3 Gbp this would sample ~13% of the genome.

**Figure 1:**
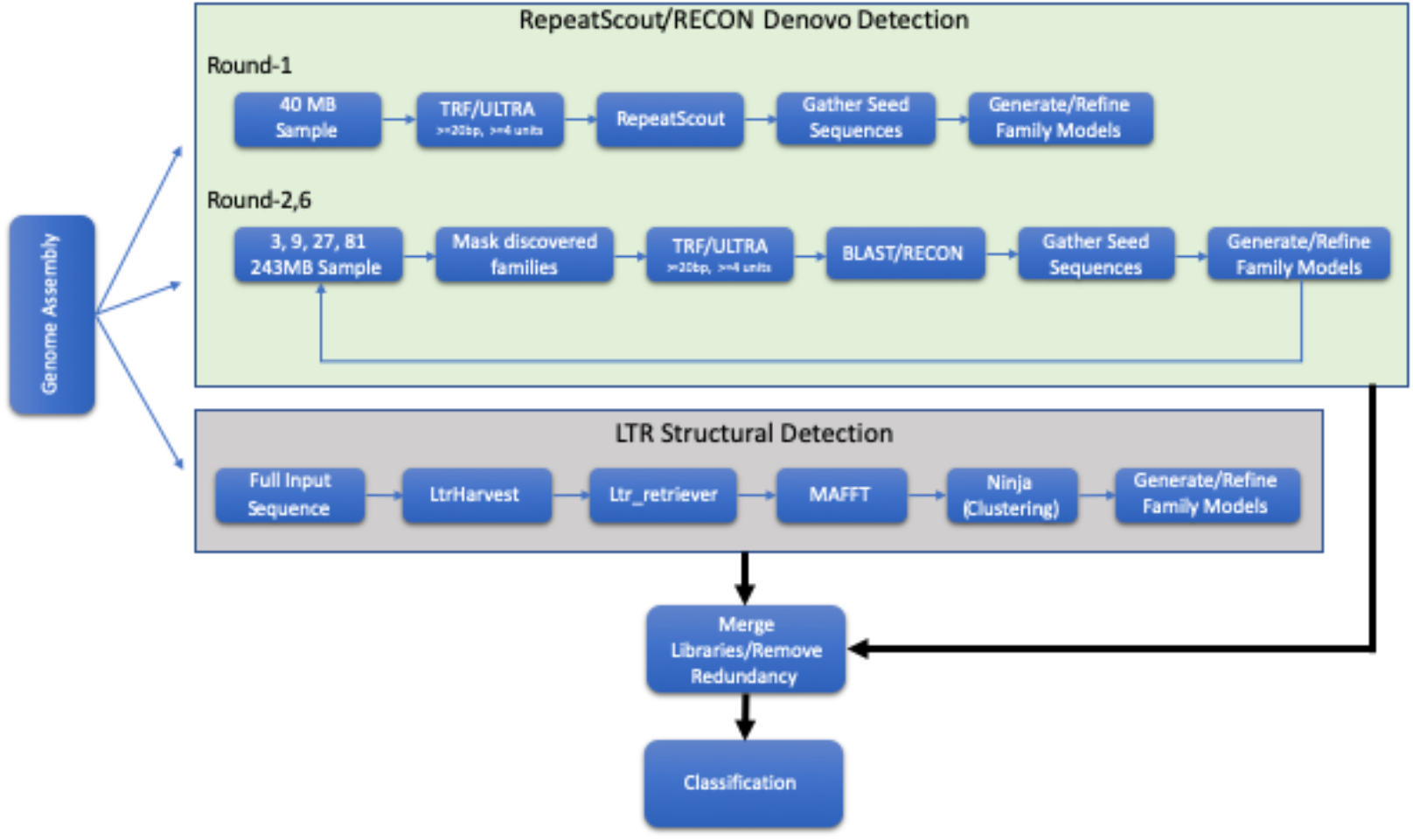
RepeatModeler2 Flow Diagram.

The new RepeatModeler2 pipeline now includes an additional structural discovery approach to assist in the discovery LTR retrotransposons. Due to their unique structure and biology (two long terminal repeat flanking a large 5-18 kb internal region) LTRs are often identified as fragments with disassociated LTR and internal regions or missing the internal segment completely using the RepeatScout/RECON methodologies. The LTR discovery is run on the complete unmasked genomic sequence and as such produces high quality LTR families with some redundancy to the previously discovered families. Therefore, we follow up LTR discovery with a merging and redundancy removal process as described in the LTR module section.

Due to constraints in the methods employed by TE discovery algorithms, their output is often either in the form of a complete or partial set of genome annotations (location ranges within the input sequence representing instances of a particular repeat), or is simply the pre-calculated consensus for each discovered TE family. In addition, the quality of these pre-calculated consensus sequences may vary considerably. A basic goal of RepeatModeler has been to produce a high-quality seed alignment and consensus sequence for each TE family; therefore we developed a seed alignment refinement method (see Refiner section below) which is employed on all families produced by the *de novo* tools.

Once all rounds of discovery/refinement/merging are complete, the final library is run through a simple classification step (see Classification section for details), where each family is assigned (if possible) to a known TE class using the unified RepeatMasker/Dfam classification nomenclature.

### Refiner description

Given a set of putatively related TE family instances the RepeatModeler Refiner tool attempts to build a high-quality seed alignment and derive from it a consensus sequence for the family. Refiner bootstraps the generation of a seed alignment by selecting as the initial consensus, the sequence which scores the best against all others. From this initial alignment a new consensus is generated and the process is repeated until the consensus stabilizes. Refiner employs an algorithm to identify low scoring subregions in the seed alignment, often caused by common indels relative to the consensus, and resolves them by globally aligning the sequences within the subregion and updating the consensus (see Supplemental Material). The consensus is not simply based on alignment column majority rule, rather each consensus position represents the highest scoring base using a scoring matrix similar to those developed for RepeatMasker, which reflects observed neutral substitution patterns in mammals. For instance, the algorithm is aware of the rapid decay of CpG sites to CpA and TpG dinucleotides in most eukaryotes due to accelerated deamination of methylated cytosines (Sved and Bird 1990; Colot and Rossignol 1999) and calls a CG pair given enough instances of aligned CA and TG dinucleotides (see supplementary data for details). The final output of the analysis is a consensus sequence and a seed alignment in Stockholm format. The latter can be used for generating profile Hidden Markov Models (Wheeler and Eddy 2013) and preserves the provenance of the family’s representative sequences.

### LTR module description

RepeatModeler2 uses the LTRharvest (Ellinghaus *et al*. 2008) package for structural LTR detection for both its overall sensitivity and speed (Ou *et al*. 2018, Ou *et al*. 2019). LTRharvest is both a discovery and annotation algorithm that does not attempt to group LTR instances into families. In addition, any region in the genome demonstrating LTR-like structure (flanking repetitive sequence of the correct size, with the correct intervening sequence) is often incorrectly identified as an LTR instance. To solve this problem, Ou and Jiang developed LTR_retriever (Ou and Jiang 2018), a package for filtering false positive results, resolving nested (mosaic) annotations, and identifying internal regions of LTRs. Some genomes have challenging nested structures that are not always resolved by LTR_retriever, so we implemented an optional parameter (-LTRMaxSeqLen) that sets the maximum allowable LTR internal length to avoid inclusion of missed mosaic internal elements in the seed alignment (see supplementary material).

We use LTR_retriever’s redundant library and perform our own clustering and consensus building process. Since LTR elements frequently contain internal deletions, and this often results in “over-splitting” elements when clustering with CD-HIT (Li and Godzik 2006), we implemented a clustering approach that scores alignment gaps as a reduced (single position) penalty. This step involves a multiple sequence alignment with MAFFT (Katoh *et al*. 2002), followed by nearest neighbor clustering into families with Ninja (Wheeler 2009). Families are then refined and consensus sequences generated in a similar fashion to results from RepeatScout/RECON.

### Combining libraries and reducing redundancy

Combining results from multiple tools is a difficult but essential step for the production of a comprehensive and non-redundant library of TE families. The RepeatScout and RECON analysis rounds reduce redundancy by masking out previously identified families with each iteration. With the addition of the LTR structural module as an independent analysis on the genome, we cannot avoid generating redundant LTR TE families. We tackled this problem by clustering the sequences between the modules with CD-HIT. Whenever RepeatModeler sequences cluster with an LTR pipeline sequence, we retain the LTR pipeline family as the representative. In addition, in RepeatModeler2 we extended this method to RepeatScout/RECON-produced families by labeling closely matching sequences as putative subfamilies (with a link to the accepted representative). Users can then choose whether to remove these subfamilies depending on the goals of their analyses.

### Classification

RepeatModeler contains a basic homology-based classification module (RepeatClassifier) which compares the TE families generated by the various *de novo* tools to both the RepeatMasker Repeat Protein DB and to the RepeatMasker libraries (e.g. Dfam and/or RepBase). The Repeat Protein DB is a set of TE-derived coding sequences that covers a wide range of TE classes and organisms. As is often the case with a search against all known TE sequences, there will be a high number of false positive or partial matches. RepeatClassifier uses a combination of score and overlap filters to produce a reduced set of high confidence results. If there is a concordance in classification among the filtered results, RepeatClassifier will label the family using the RepeatMasker/Dfam classification system and adjust the orientation (if necessary). Remaining families are labeled “Unknown” if a call cannot be made.

### Benchmarking

We benchmarked RepeatModeler2 on model species that have high-quality reference TE libraries: *D. melanogaster, D. rerio*, and *O. sativa*. We used Repbase for the *D. melanogaster* (release 20181026) and *D. rerio* (release 14.01) libraries, and the manually-improved library for *O. sativa* from (Ou *et al*. 2019). We used RepeatMasker and parseRM (https://github.com/4ureliek/Parsing-RepeatMasker-Outputs/blob/master/parseRM.pl) to estimate the percentage of the genome masked by each subclass for the manually-curated and RepeatModeler2 libraries.

An important aspect of our pipeline is its ability to produce accurate consensus sequences corresponding to unique TE families. Thus, we also assessed the quality of our families by comparing their sequences with the sequences of the manually-curated consensus sequences of reference libraries. We used RepeatMasker (v 1.332) with the reference library, and the RepeatModeler2 output library as the subject. We then used a custom bash script (https://github.com/jmf422/TE_annotation/blob/master/get_family_summary_paper.sh) to assess the sequence identity and coverage of matches between the libraries. We classify families as being “perfect”, “good”, “present”, or “not found” based on the following definitions. “Perfect” families are those for which one sequence in our *de novo* library matches with >95% nucleotide similarity and >95% length coverage to a family consensus in the reference library. “Good” families are those in which multiple overlapping sequences in our output library match with >95% nucleotide similarity and >95% coverage to the curated consensus. A family is considered “present” if one or multiple library sequences align with >80% similarity and >80% coverage to the reference consensus sequence. Below these thresholds, we consider a family “not found” (although there may be fragments present in our output library).

## Results and Discussion

We benchmarked RepeatModeler2 on three model species (fruit fly, zebrafish, rice) with diverse TE landscapes for which reference TE libraries have been extensively curated over decades of study (Figure 2). As a first assessment of the ability of RepeatModeler2 to capture known TEs in each of these genomes, we use each output library to run RepeatMasker against the cognate genome assembly and measured the percent of the genome masked by each major TE subclass. We also counted the number of sequences in each library falling within each subclass. RepeatModeler2 was able to recover accurately the contrasted TE landscapes of these species (compare with curated libraries in Figure 2).

**Figure2:**
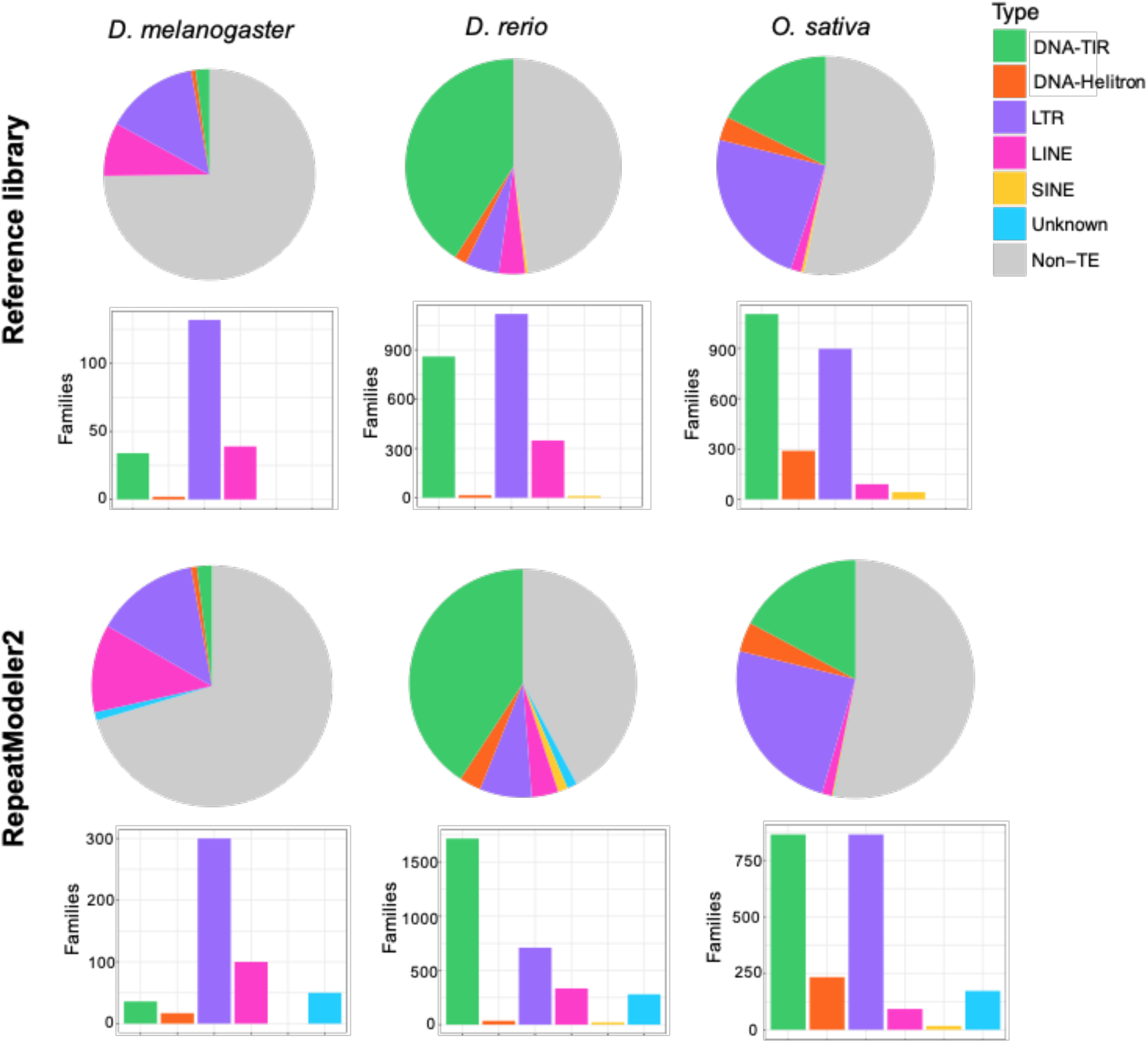
Benchmarking of RepeatModeler2 on three model species. Top panel: genome composition (top) and number of families (bottom) of each TE subclass for the reference libraries. Bottom panel: genome composition (top) and number of families (bottom) of each TE subclass for the RepeatModeler2 library.

As previously documented (e.g. Lerat et al. 2003), our results show that the genome of the fruit fly *D. melanogaster* is dominated by retrotransposons, especially LTR retroelements. This is reflected both by the amount of genomic DNA covered by these elements and by the number of unique families (Figure 2). The zebrafish, *D. rerio*, is dominated by class II, DNA-TIR transposons, but also exhibits a very diverse assortment of LTR retroelements with many unique families (Howe *et al*. 2013). While our RepeatModeler2 library captures this general composition, it defines about twice less LTR families but twice more TIR families than in the original reference library. These differences may be caused by 1) a less stringent definition of LTR families in the reference library compared to the RM2 library; and 2) the fact that DNA-TIR elements are only identified by the RECON/RepeatScout module of RepeatModeler2, which tend to produce shorter, more fragmented sequences than the LTR structural module. Therefore the number of sequences classified as DNA-TIRs by RepeatModeler2 may be inflated and may rather represent variants or fragments of the same family (see Figure 3). The genome of rice, *O. sativa*, is known to contain almost equal proportions and numbers of DNA-TIR and LTR elements (Ou *et al*. 2019), and this profile is recovered by our RepeatModeler2 library (Figure 2). In summary, RepeatModeler2 produces libraries that recapitulate the major TE subclass composition of these three model species.

**Figure 3:**
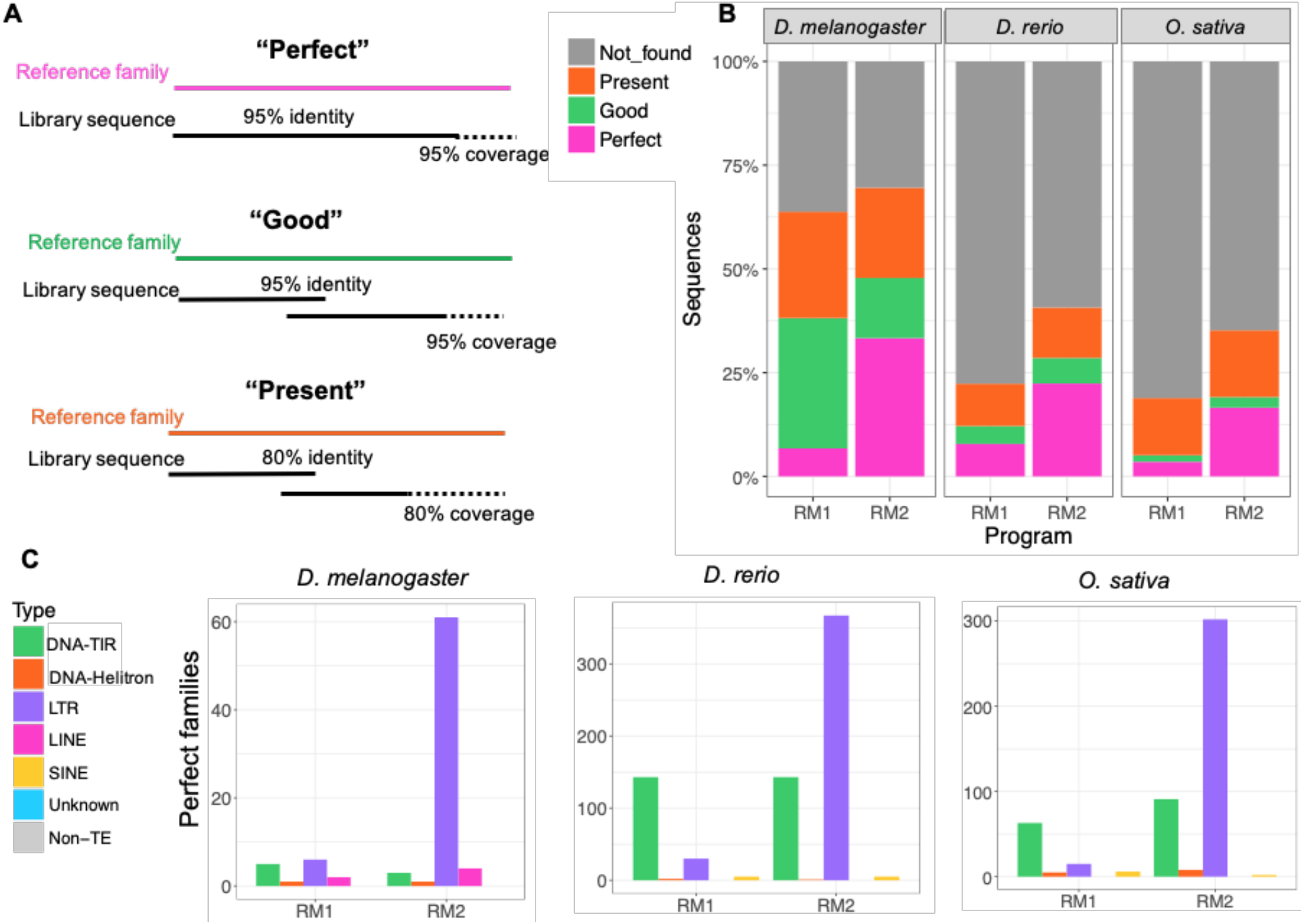
Evaluation family by family for RepeatModeler1 and RepeatModeler2. (A) Definitions of “Perfect”, “Good”, and “Present” families. “Perfect” families are those for which one sequence in our *de novo* library matches >95% in sequence identity and coverage to a family in the reference library. “Good” families are those in which multiple overlapping library sequences with alignments >95% similar to the reference consensus make up the >95% sequence coverage of the element. Finally, a family is considered “present” if one or multiple library sequences align with >80% similarity to the reference consensus sequence and cover >80% of the sequence. Otherwise, we consider a family “not found” (although there may be fragments present) (B) Summary of families found by the last release of RepeatModeler (RM1) and RepeatModeler2 (RM2). (C) Number of perfect families by subclass for each benchmark species.

Next we assess the ability of RepeatModeler2 to accurately capture the diversity and sequence of unique TE families. RepeatModeler2 produced 766, 3851, and 2648 library sequences for *D. melanogaster, D. rerio*, and *O.sativa*, respectively - all comparable to the number of individual sequences in the reference libraries (Table 1). In addition, RepeatModeler2 labels families that cluster within 20% similarity as “putative subfamily”, thus we also provided the number of sequences without the inclusion of subfamilies (Table 1). A TE library with more sequences is not necessarily more useful since it often indicates redundancy and fragmented sequences. Since we use a redundancy-removal step, RepeatModeler2 did not produce drastically more family models than the previous version of the program.

**Table 1:** Number of sequences produced from RepeatModeler and present in the manually-curated libraries. The second column indicates the total number of families produced by RepeatModeler2 and the third column is the same except not including those annotated as subfamilies. The fourth column is the number of sequences in the curated library, and the fifth column is the number of sequences produced from RepeatModeler1.

| Species | RM2 Families | RM2 Families (no subfamilies) | Curated library | RM1 families |
| --- | --- | --- | --- | --- |
| *D. melanogaster* | 734 | 509 | 207 | 699 |
| *D. rerio* | 3851 | 3147 | 2322 | 2503 |
| *O. sativa* | 2648 | 2284 | 2431 | 1652 |

The most significant improvement of RepeatModeler2 over the previous release of the program is in the quality (accuracy) of the family consensus sequences delivered in the output library (Figure 3). We labelled the sequence matches between the RepeatModeler2 and reference libraries as “perfect”, “good”, or “present” based on the level of sequence similarity and coverage (Figure 3A, see also Methods). If the family did not meet the minimum criteria for being counted as “present”, it was reported as “not found”. RepeatModeler2 produced 2.9 to 4.7 times more “perfect” families than the first version of RepeatModeler, and had more sequences closely matching the reference libraries overall (Figure 3B). Most of this improvement can be attributed to perfect LTR element families that are identified by RepeatModeler2, but were previously missed (Figure 3C). The evaluation criteria we used for families were relatively strict, probably explaining the large number of families “not found”. Indeed, when masking each genome with the cognate library, we were able to recapitulate the genomic proportions of each subclass as generally obtained with the reference library (Figure 2).

Eukaryotic genomes contain complex structure and tandem repeats, which may result in false positives for TE discovery software. We assessed the false positive rate of RepeatModeler2 by running it on artificially-generated genomes devoid of TEs simulated by GARLIC (Caballero *et al*. 2014) for *D. melanogaster* and *D. rerio*. GARLIC generates background sequences with realistic complexity, isochore structure and tandem repeat content as the modeled genome. RepeatModeler2 produced only one false positive family on the *D. melanogaster* artificial genome and five false positive families on the *D. rerio* artificial genome. No false positive families were produced from the LTR module. These results suggest that the rate of false positives generated by RepeatModeler2 is very low.

In addition to LTR retroelements, other types of TEs are sometimes considered challenging to identify *ab initio* with automated approaches. In particular, MITEs (miniature inverted-repeat transposable elements) can be challenging because they are typically non-coding and may be highly diverged in sequence from from their parental DNA:TIR element (Feschotte *et al*. 2002; Han and Wessler 2010). The zebrafish and rice genomes are known to contain a large number of MITE families (Jiang *et al*. 2004), but these elements did not appear under-represented in the RepeatModeler2 libraries generated for these two species (Figure 2). In comparison to the reference libraries, RepeatModeler2 also performed well with the identification of Helitrons, which also pose particular challenges for automated discovery (reviewed in (Thomas and Pritham 2015)) (Figure 2). Thus, RepeatModeler2 appears capable of recovering a wide diversity of TEs which have been traditionally considered recalcitrant to ab initio identification.

We anticipate that additional improvements will further enhance the current version of the pipeline. Because of the modular architecture of RepeatModeler2, it should be relatively straightforward to add other modules tailored to the discovery of specific subclass of elements such as those dedicated to the identification of MITEs (Han and Wessler 2010) or Helitrons (Yang and Bennetzen 2009; Xiong *et al*. 2014). It is also important to emphasize that the classifier currently used by RepeatModeler2 remains rudimentary as it strictly relies on detection of sequence homology to known TEs and protein domains. This limitation hampers the ability to classify non-coding elements or elements with sequences highly diverged from those annotated in the databases. Integrating additional features used for TE classification, such as conserved TIR sequence motifs or target site duplications, as implemented in other TE classifiers (Feschotte *et al*. 2009) will further improve the ability of RepeatModeler2 to deliver high-quality libraries.

The genome annotation community is in pressing need of a TE discovery program that is easy to use, has been thoroughly benchmarked, and can be applied to almost any eukaryotic species (Hoen *et al*. 2015). We believe that RepeatModeler2 will meet this demand. Other TE discovery programs exist, but either focus on finding instances in the genome instead of family consensus sequence building (e.g. Berthelier *et al*. 2018, Ou *et al*. 2019), or are challenging to install and use (Flutre *et al*. 2011). RepeatModeler2 is easy to install and run, as we provide a container version to avoid installing independently all dependencies. We anticipate that the application of RepeatModeler2 to existing and future genome assemblies will result in more consistent genome annotations and improved TE family models, which will enhance a wide array of genomic analyses including but not limited to TE biology.

## Acknowledgements

We thank Arnie Kas, Warren Gish, Alkes Price, Pavel Pevzner, Shujun Ou, and Ning Jiang for assistance with dependencies used by RepeatModeler. We thank Andy Siegel for statistics consultations in the development of RepeatModeler. This work was supported by NIH grants U01-HG009391 and R35-GM122550 to CF, and NHGRI grant U24 HG010136 and NHGRI R01 HG002939 to AFS. JMF was supported by a NSERC PGSD graduate fellowship.

## SUPPLEMENTARY MATERIALS

### LTRPipeline

The development of RepeatModeler2 was motivated by author JMF working on annotating transposable elements in Drosophila with CG, AGC, and CF. In Drosophila, LTR elements are the most abundant subclass of TEs and are often present in heterochromatic regions where they frequently contain nested structures. JMF developed a bash pipeline incorporating RepeatModeler and an LTR-specific module, which resulted in greatly improved TE libraries for Drosophila. We found that this pipeline also performed better than RepeatModeler in other species we tested it on such as zebrafish. We believed that the TE community at large could benefit from the incorporation of the LTR pipeline, so we partnered with the authors of RepeatModeler (RH, JR, and AFS) to incorporate the LTR pipeline into an improved software we call RepeatModeler2.

One of the main troubleshooting issues with incorporating a structural LTR identification program was that it identifies instances of LTR elements in the genome, whereas the goal of the RepeatModeler is to identify TE families. In order to remove redundancy and have one seed alignment (and consensus) per family, sequence similarity clustering is used. LTR_retriever performs similarity clustering with CD-HIT to accomplish this. However, we noticed that the CD-HIT clustered output of LTR_retriever still contained redundancy. We hypothesized that this was because of CD-HIT’s gap scoring policy, as it scores each base pair of an indel as a penalty, rather than the indel as a single penalty, which is more biologically relevant (Flynn *et al*. 2015). This issue is especially evident for LTR elements, which commonly contain internal deletions. We originally used a multiple sequence alignment with MAFFT followed by nearest-neighbor clustering with MOTHUR (Schloss *et al*. 2009). MOTHUR clustering scores indels as a single gap penalty, and we found it worked to effectively cluster LTR elements into accurate family groups. In later versions of the pipeline, we made a custom clustering script with NINJA using the same procedure as MOTHUR. Instead of using the longest sequence in the cluster as the family representative as LTR_retriever does, we incorporated the Refiner module from RepeatModeler to build seed alignments and a family consensus from the cluster members.

### LTR internal length filtering

LTR elements often acquire nested insertions. In RepeatModeler2 we utilize the LTR_retriever analysis tool, which attempts to remove nested insertions; however we found that some nested insertions still remain. This can reduce the quality of the LTR library. One way to prevent LTR elements with nested insertions from being included in the LTR library is to impose a maximum length of the internal sequence (optional parameter −LTRMaxSeqLen). For example, in Drosophila, almost all true LTR internal elements are <10 kb and it is known that nested LTR elements are common in heterochromatic regions, we would use −LTRMaxSeqLen 10000. Since using this parameter could potentially bias the results, we only recommend using it if the max LTR length is known or if nested insertions are a problem in the particular analysis.

**Figure S1:**
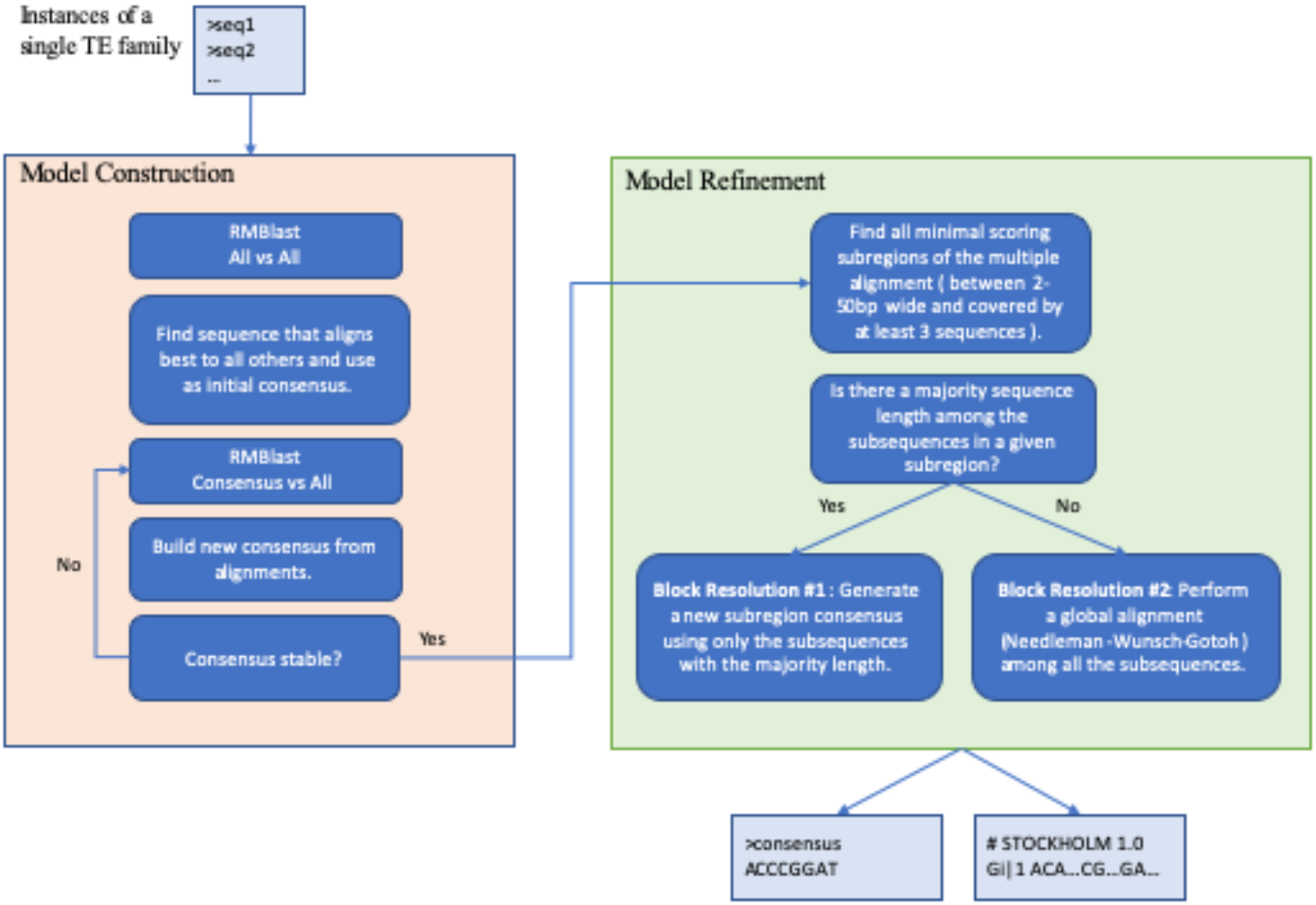
RepeatModeler Refiner module flowchart.

### Family Refinement

The RepeatModeler Refiner tool produces a multiple sequence alignment (seed alignment) from putatively related instances of a TE family. Refiner uses an iterative process (Figure S1) to build and then improve the seed alignment for the family. This process is bootstrapped by first performing a full pairwise comparison of all instance sequences to each other and selecting, as the initial consensus, the instance which aligns best to all others. It is possible that the initial sequence chosen does not align to a small portion of the input sequences. In this case the unaligned sequences are maintained in a candidate pool for possible inclusion in further iterations of consensus refinement. The pairwise alignments to the chosen sequence are combined into the initial seed alignment and novel CpG-adjusted consensus caller is used to generate an updated consensus for the family.

The consensus caller used in RepeatModeler differs from a standard majority-rule consensus caller in two ways: it scores each possible ancestral base or IUB code to the seed alignment column using a neutral substitution matrix, and it looks for overrepresentation of common CpG mutation products to correctly identify the ancestral state of CpG dimers. The first step uses a matrix (Figure S2) that reflects observed neutral DNA substitution patterns. This matrix and similar matrices used by RepeatMasker were derived from studies of neutrally decaying DNA transposon families in mammals.

**Figure S2:**
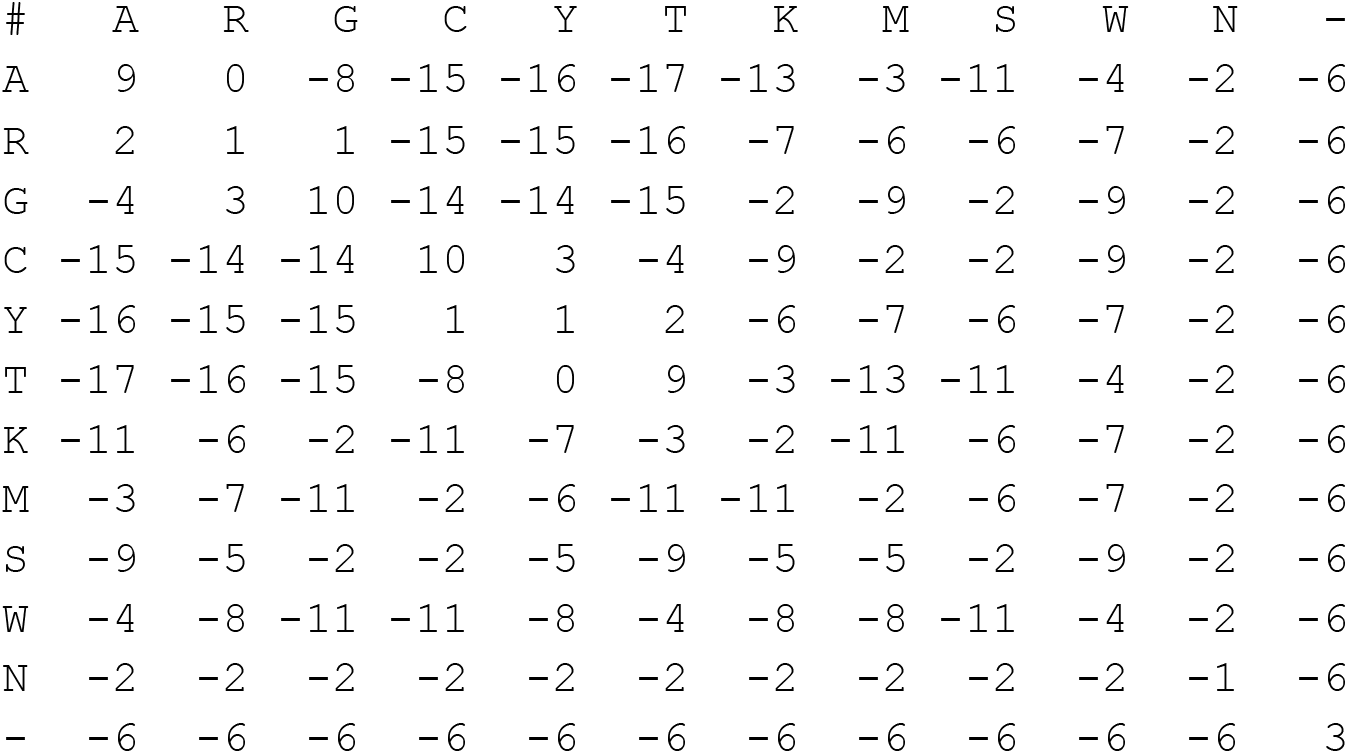
Neutral substitution matrix used by the RepeatModeler consensus caller. Rows and columns represent the ancestral and current nucleic acids respectively. IUPAC codes, as well as a standard gap (“-”) characters are included.

After the score for each possible ancestral base and at each column in the alignment is calculated, a 2 bp sliding window is run over the alignment and the highest dimer score is calculated for window. Using the truth table (Table S3), an alternative score is developed for the hypothesis that the ancestral dimer was a CpG (CG_Score). If the total CG_Score is higher than the matrix score for the window the consensus is changed to a “CG” at these two positions in the alignment.

**Table S3:** Dimer score table for calling CpG sites. This table represents the dimers which are most likely to be generated by mutation at CpG site. For these cases a modified matrix score (CG Score) is used to calculate the alternative score for the alignment columns.

| Observation | Matrix Score | CG Score | Description |
| --- | --- | --- | --- |
| TG | +6 | +12 | Direct result of current strand CpG deaminating the C and converting to a TG. |
| CA | +6 | +12 | Indirect result of CpG on opposite strand converting to TG and an incorrect repair of the current strand. |
| TA | -8 | -5 | Two step result. CG -> TG followed by a common transition of either forward strand TG -> TA or reverse strand CA -> TA. |
| TT | -19 | -13 | CpG transition + transversion. E.g. CG -> TG -> TT. |
| TC | -18 | -13 | (dito) CG -> TG -> TC. |
| AA | -19 | -13 | (dito) CG -> CA -> AA. |
| GA | -18 | -13 | (dito) CG -> CA -> GA. |

The updated consensus is re-aligned to all seed alignment sequences as well as the candidate pool and the process is repeated until the consensus sequence stabilizes or a maximum iteration count is exceeded. At this stage the seed alignment may often contain short regions of misalignment caused by tandem duplications or deletions within the original consensus choice. We identify these regions by calculating seed alignment quality using a sliding window approach (Ruzzo and Tompa 1999) and considering each region independently for consensus refinement. If there is a majority sequence length within the region, the consensus is called from only the sequences of this length, otherwise an all-vs-all global alignment is performed and the sequence scoring best against all others is used to align and then call the new sub-region consensus. The sub-regions consensi are replaced in the full-consensus and the original consensus refinement process is repeated until the consensus stabilizes.

### Benchmarking Parameters

The benchmarking analysis was performed on a research group-shared CentOS 7.6.1810 Linux machine (Intel(R) Xeon(R) CPU E7-4830 v4 @ 2.00GHz, 112 cores, 504 GB RAM). RepeatModeler utilizes a seeded random number generator in the selection of the genomic samples. The seed is automatically chosen and displayed at runtime to facilitate reproducibility (using the −srand option). The runtime, and seeds are shown in Table S2. RepeatModeler 1.0.11 benchmarks were generated using only the “−database” parameter using the program defaults. The new “−LTRStruct” option to RepeatModeler2 was used in all benchmark runs. In addition for *D. melanogaster*, the −LTRMaxSeqLen 10000 was used.

**Table S2:** benchmarking information. RM = RepeatModeler1.0.11, RM2 = RepeatModeler 2.

| Species | Assembly size (Mb) | RM Seed # | RM cores | RM runtime | RM2 Seed # | RM2 cores | RM2 runtime |
| --- | --- | --- | --- | --- | --- | --- | --- |
| *D. melanogaster* | 164 | 1572903982 | 8 | 62:26:16 | 1570222393 | 8 | 12:55:59 |
| *D. rerio* | 1372 | 1570465657 | 16 | 33:12:10 | 1570222833 | 16 | 40:35:56 |
| *O. sativa* | 375 | 1570476015 | 16 | 81:03:54 | 1570222971 | 16 | 37:22:46 |

Genome assembly versions used for benchmarking:

*D. melanogaster* - PBcr-BLASR Celera 8.1 assembly (Koren et al. 2012)
*D. rerio* - danRer10: https://www.ncbi.nlm.nih.gov/assembly/GCF_000002035.5/
*O. sativa* - Verson 7.0 http://rice.plantbiology.msu.edu/pub/data/Eukaryotic_Projects/o_sativa/annotation_dbs/pseudomolecules/version_7.0/all.dir/all.con

The scripts used to produce the benchmarking statistics may be found here: (https://github.com/Dfam-consortium/RepeatModeler). RepeatModeler2 was configured using the following versions of dependent packages:

- Mafft 7.407
- RepeatMasker 4.0.9-p2
- RECON 1.0.8
- RepeatScout 1.0.6
- RMBLast 2.9.0-p2
- TRF 4.0.9
- Ninja 0.97-cluster_only
- CD-HIT 4.8.1
- GenomeTools 1.5.10
- LTR_retriever 2.6

## REFERENCES

Arkhipova I. R., 2017 Using bioinformatic and phylogenetic approaches to classify transposable elements and understand their complex evolutionary histories. Mob. DNA 8: 19.

Bao Z., 2002 Automated De Novo Identification of Repeat Sequence Families in Sequenced Genomes. Genome Research 12: 1269–1276.

Bao W., K. K. Kojima, and O. Kohany, 2015 Repbase Update, a database of repetitive elements in eukaryotic genomes. Mob. DNA 6: 11.

Berthelier J., N. Casse, N. Daccord, V. Jamilloux, B. Saint-Jean, et al., 2018 A transposable element annotation pipeline and expression analysis reveal potentially active elements in the microalga Tisochrysis lutea. BMC Genomics 19: 378.

Bourque G., K. H. Burns, M. Gehring, V. Gorbunova, A. Seluanov, et al., 2018 Ten things you should know about transposable elements. Genome Biol. 19: 199.

Caballero J., A. F. A. Smit, L. Hood, and G. Glusman, 2014 Realistic artificial DNA sequences as negative controls for computational genomics. Nucleic Acids Res. 42: e99.

Chang C.-H., and A. M. Larracuente, 2019 Heterochromatin-Enriched Assemblies Reveal the Sequence and Organization of the Y Chromosome. Genetics 211: 333–348.

Colot V., and J. L. Rossignol, 1999 Eukaryotic DNA methylation as an evolutionary device. Bioessays 21: 402–411.

Consortium I. H. G. S., and International Human Genome Sequencing Consortium, 2001 Initial sequencing and analysis of the human genome. Nature 409: 860–921.

Ellinghaus D., S. Kurtz, and U. Willhoeft, 2008 LTRharvest, an efficient and flexible software for de novo detection of LTR retrotransposons. BMC Bioinformatics 9: 18.

Feschotte C., S. R. Wessler, and X. Zhang, 2002 Miniature Inverted-Repeat Transposable Elements and Their Relationship to Established DNA Transposons. Mobile DNA II 1147–1158.

Feschotte C., and E. J. Pritham, 2007 DNA transposons and the evolution of eukaryotic genomes. Annu. Rev. Genet. 41: 331–368.

Feschotte C., U. Keswani, N. Ranganathan, M. L. Guibotsy, and D. Levine, 2009 Exploring repetitive DNA landscapes using REPCLASS, a tool that automates the classification of transposable elements in eukaryotic genomes. Genome Biol. Evol. 1: 205–220.

Finnegan D. J., 1989 Eukaryotic transposable elements and genome evolution. Trends Genet. 5: 103–107.

Flutre T., E. Duprat, C. Feuillet, and H. Quesneville, 2011 Considering transposable element diversification in de novo annotation approaches. PLoS One 6: e16526.

Grabundzija I., A. B. Hickman, and F. Dyda, 2018 Helraiser intermediates provide insight into the mechanism of eukaryotic replicative transposition. Nat. Commun. 9: 1278.

Han Y., and S. R. Wessler, 2010 MITE-Hunter: a program for discovering miniature inverted-repeat transposable elements from genomic sequences. Nucleic Acids Res. 38: e199.

Hoen D. R., G. Hickey, G. Bourque, J. Casacuberta, R. Cordaux, et al., 2015 A call for benchmarking transposable element annotation methods. Mobile DNA 6, 13.

Howe K., M. D. Clark, C. F. Torroja, J. Torrance, C. Berthelot, et al., 2013 The zebrafish reference genome sequence and its relationship to the human genome. Nature 496: 498–503.

Huang C. R. L., K. H. Burns, and J. D. Boeke, 2012 Active transposition in genomes. Annu. Rev. Genet. 46: 651–675.

Hua-Van A., A. Le Rouzic, C. Maisonhaute, and P. Capy, 2005 Abundance, distribution and dynamics of retrotransposable elements and transposons: similarities and differences. Cytogenet. Genome Res. 110: 426–440.

Hubley R., R. D. Finn, J. Clements, S. R. Eddy, T. A. Jones, et al., 2016 The Dfam database of repetitive DNA families. Nucleic Acids Res. 44: D81–9.

International Wheat Genome Sequencing Consortium (IWGSC), IWGSC RefSeq principal investigators:, R. Appels, K. Eversole, C. Feuillet, et al., 2018 Shifting the limits in wheat research and breeding using a fully annotated reference genome. Science 361. https://doi.org/10.1126/science.aar7191

Jiang N., C. Feschotte, X. Zhang, and S. R. Wessler, 2004 Using rice to understand the origin and amplification of miniature inverted repeat transposable elements (MITEs). Curr. Opin. Plant Biol. 7: 115–119.

Jiao Y., P. Peluso, J. Shi, T. Liang, M. C. Stitzer, et al., 2017 Improved maize reference genome with single-molecule technologies. Nature 546: 524–527.

Jurka J., V. V. Kapitonov, O. Kohany, and M. V. Jurka, 2007 Repetitive Sequences in Complex Genomes: Structure and Evolution. Annual Review of Genomics and Human Genetics 8: 241–259.

Katoh K., K. Misawa, K.-I. Kuma, and T. Miyata, 2002 MAFFT: a novel method for rapid multiple sequence alignment based on fast Fourier transform. Nucleic Acids Res. 30: 3059–3066.

Koepfli K.-P., B. Paten, Genome 10K Community of Scientists, and S. J. O’Brien, 2015 The Genome 10K Project: a way forward. Annu Rev Anim Biosci 3: 57–111.

Lerat E., C. Rizzon, and C. Biémont, 2003 Sequence divergence within transposable element families in the Drosophila melanogaster genome. Genome Res. 13: 1889–1896.

Lewin H. A., G. E. Robinson, W. J. Kress, W. J. Baker, J. Coddington, et al., 2018 Earth BioGenome Project: Sequencing life for the future of life. Proc. Natl. Acad. Sci. U. S. A. 115: 4325–4333.

Li W., and A. Godzik, 2006 Cd-hit: a fast program for clustering and comparing large sets of protein or nucleotide sequences. Bioinformatics 22: 1658–1659.

Ou S., J. Chen, and N. Jiang, 2018 Assessing genome assembly quality using the LTR Assembly Index (LAI). Nucleic Acids Research. 46:e126

Ou S., and N. Jiang, 2018 LTR_retriever: A Highly Accurate and Sensitive Program for Identification of Long Terminal Repeat Retrotransposons. Plant Physiology 176: 1410–1422.

Ou S., W. Su, Y. Liao, K. Chougule, D. Ware, et al. 2019 Benchmarking Transposable Element Annotation Methods for Creation of a Streamlined, Comprehensive Pipeline. Biorxiv https://doi.org/10.1101/657890

Piégu B., S. Bire, P. Arensburger, and Y. Bigot, 2015 A survey of transposable element classification systems--a call for a fundamental update to meet the challenge of their diversity and complexity. Mol. Phylogenet. Evol. 86: 90–109.

Price A. L., N. C. Jones, and P. A. Pevzner, 2005 De novo identification of repeat families in large genomes. Bioinformatics 21 Suppl 1: i351–8.

Saha S., S. Bridges, Z. V. Magbanua, and D. G. Peterson, 2008a Computational Approaches and Tools Used in Identification of Dispersed Repetitive DNA Sequences. Tropical Plant Biology 1: 85–96.

Saha S., S. Bridges, Z. V. Magbanua, and D. G. Peterson, 2008b Empirical comparison of ab initio repeat finding programs. Nucleic Acids Res. 36: 2284–2294.

Smit A. F., 1999 Interspersed repeats and other mementos of transposable elements in mammalian genomes. Curr. Opin. Genet. Dev. 9: 657–663.

Smit, Arian. “RepeatMasker Genomic Datasets.” http://www.repeatmasker.org/genomicDatasets/RMGenomicDatasets.html, 22 Mar. 2012. Web

Stitzer M. C., S. N. Anderson, N. M. Springer, and J. Ross-Ibarra, 2019 The Genomic Ecosystem of Transposable Elements in Maize. Biorxiv https://doi.org/10/1101/559922

Sved J., and A. Bird, 1990 The expected equilibrium of the CpG dinucleotide in vertebrate genomes under a mutation model. Proc. Natl. Acad. Sci. U. S. A. 87: 4692–4696.

Thomas J., and E. J. Pritham, 2015 Helitrons, the Eukaryotic Rolling-circle Transposable Elements. Microbiol Spectr 3. https://doi.org/10.1128/microbiolspec.MDNA3-0049-2014

Vargiu L., P. Rodriguez-Tomé, G. O. Sperber, M. Cadeddu, N. Grandi, et al., 2016 Classification and characterization of human endogenous retroviruses; mosaic forms are common. Retrovirology 13: 7.

Wheeler T. J., 2009 Large-Scale Neighbor-Joining with NINJA. Lecture Notes in Computer Science 375–389.

Wheeler T. J., and S. R. Eddy, 2013 nhmmer: DNA homology search with profile HMMs. Bioinformatics 29: 2487–2489.

Wicker T., F. Sabot, A. Hua-Van, J. L. Bennetzen, P. Capy, et al., 2007 A unified classification system for eukaryotic transposable elements. Nat. Rev. Genet. 8: 973–982.

Xiong W., L. He, J. Lai, H. K. Dooner, and C. Du, 2014 HelitronScanner uncovers a large overlooked cache of Helitron transposons in many plant genomes. Proc. Natl. Acad. Sci. U. S. A. 111: 10263–10268.

Yandell M., and D. Ence, 2012 A beginner’s guide to eukaryotic genome annotation. Nat. Rev. Genet. 13: 329–342.

Yang L., and J. L. Bennetzen, 2009 Structure-based discovery and description of plant and animal Helitrons. Proc. Natl. Acad. Sci. U. S. A. 106: 12832–12837.

## References

Flynn J. M., E. A. Brown, F. J. J. Chain, H. J. MacIsaac, and M. E. Cristescu, 2015 Toward accurate molecular identification of species in complex environmental samples: testing the performance of sequence filtering and clustering methods. Ecology and Evolution 5: 2252–2266.

Ruzzo W. L., and M. Tompa, 1999 A linear time algorithm for finding all maximal scoring subsequences. Proc. Int. Conf. Intell. Syst. Mol. Biol. 234–241.

Schloss P. D., S. L. Westcott, T. Ryabin, J. R. Hall, M. Hartmann, et al., 2009 Introducing mothur: open-source, platform-independent, community-supported software for describing and comparing microbial communities. Appl. Environ. Microbiol. 75: 7537–7541.

